# Low dimensional morphospace of topological motifs in human fMRI brain networks

**DOI:** 10.1101/153320

**Authors:** Sarah E. Morgan, Sophie Achard, Maite Termenon, Petra E. Vértes, Edward T. Bullmore

## Abstract

We present a low-dimensional morphospace of fMRI brain networks, where axes are defined in a data-driven manner based on the network motifs. The morphospace allows us to identify the key variations in healthy fMRI networks in terms of their underlying motifs and we observe that two principal components (PCs) can account for 97% of the motif variability. The first PC corresponds to the small-world axis and correlates strongly with the networks’ global efficiency. There is also some evidence that PC1 correlates with the average length of the 5% of longest edges in the network. Hence this axis represents the trade-off between the cost of long distance edges and their topological benefits. The second PC correlates with the networks’ assortativity. Finally, we show that the economical clustering generative model proposed by Vértes et al. can approximately reproduce the motif PC space of the real fMRI brain networks, in contrast to other generative models. Overall, the motif morphospace provides a powerful way to visualise the relationships between network properties and to study the driving forces behind the topology of fMRI brain networks.

## Introduction

Understanding the factors influencing the global topology of fMRI brain networks is a long-held ambition of network neuroscience. The energetic cost of forming and maintaining long-distance connections has been established as an important factor shaping brain networks [6], however it is also well-known that cost minimisation alone cannot explain all observed network features [15]. In 2012, Vértes et al. showed that several properties of fMRI networks can be reproduced by a generative model which encodes a trade-off between the cost of long-distance connections and the topological features they enable [30]. Important questions remain, for example which are the key network metrics that a generative model should capture and what are the relationships between different topological features of fMRI networks?

Generative models typically focus on a number of global metrics to characterise the similarity between the observed and modelled network topology, e.g. global efficiency or modularity [24, 16]. However, the plethora of metrics available can make results difficult to compare across studies, metrics often overlap [17] and a top-down approach risks missing functionally important features. We present a data driven approach using motifs to characterise brain networks, removing the need for arbitrarily chosen global metrics. Motifs are the ‘building blocks’ of networks and are known to vary between networks with different topologies [19, 18]. They have also been used to classify networks into superfamilies, with different motif profiles shaped by different functional roles [18]. In 2004, Sporns and Kötter showed that motifs can also give insight into the structure and function of brain networks [26].

After characterising our networks in terms of their motif fingerprints, we perform a principal component analysis (PCA) and find that two principal components (PCs) can explain 97% of the motif variability. This allows us to build a low-dimensional morphospace defined by these first two PCs. Morphospaces have already been shown as a promising approach to study network topology [2, 9]. To our knowledge, we present the first morphospace whose axes are based on a principal component analysis of motif fingerprints.

Our results show that the global topological properties of fMRI brain networks can be described by a low-dimensional space and that the fundamental dimensions relate to certain global network metrics. The first motif PC correlates with global efficiency. There is also some evidence that PC1 correlates with the average Euclidean distance of the longest 5% of edges, suggesting that the spatial embedding of fMRI brain networks is reflected in their global topology. The second PC correlates with the networks’ assortativity. The low-dimensional space described here agrees with prior results suggesting that very few parameters are needed to reproduce key features of the networks [30, 5, 10, 13, 14, 16]. Finally, we show that the economical clustering model proposed by [30] can largely reproduce the motif fingerprints and morphospace obtained experimentally. Interestingly, other plausible models (e.g. the economical preferential attachment model) are unable to do this, hence our motif morphospace provides a simple and powerful way to discriminate between models and evaluate model-data fit.

## Results

### Generation of the morphospace from motif fingerprints

We begin by calculating undirected fMRI networks for 100 healthy subjects from the Human Connectome Project by correlating the regional fMRI wavelet time series and thresholding the resulting correlation matrices. Details of the data and the pre-processing are given in the Methods section. Unless otherwise stated, we take a threshold of 800 edges (a connection density of ≈ 20%). We then characterise the networks in terms of their motifs. G′ = (V′, E′) is a motif of the graph G = (V, E) with vertices V and edges E, if V′ ⊆ V and E’ is a subset of E such that the vertices of each edge in E’ are in V’. In this work we consider induced motifs, for which E’ includes all edges of G which end on vertices V’ [7]. We focus on the six possible 4-node, undirected motifs, which are shown in Fig. 1a. Following [26], these motifs can be denoted as 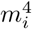, where *i* ∈ 1, 6. Since we only consider 4-node motifs, we drop the superscript 4 for brevity. We note that the different motifs have different numbers of edges and closed triangles; *m*_6_ is fully-connected, whilst *m*_1_ and *m*_3_ have the least edges and no closed triangles. Different motifs also exhibit different levels of degree heterogeneity, as shown by the colour of the nodes in Fig. 1a. *m*_3_ and *m*_6_ have nodes with the most homogeneous degrees, whilst *m*_2_ and *m*_4_ have nodes with the most heterogeneous degrees.

**Figure 1.**
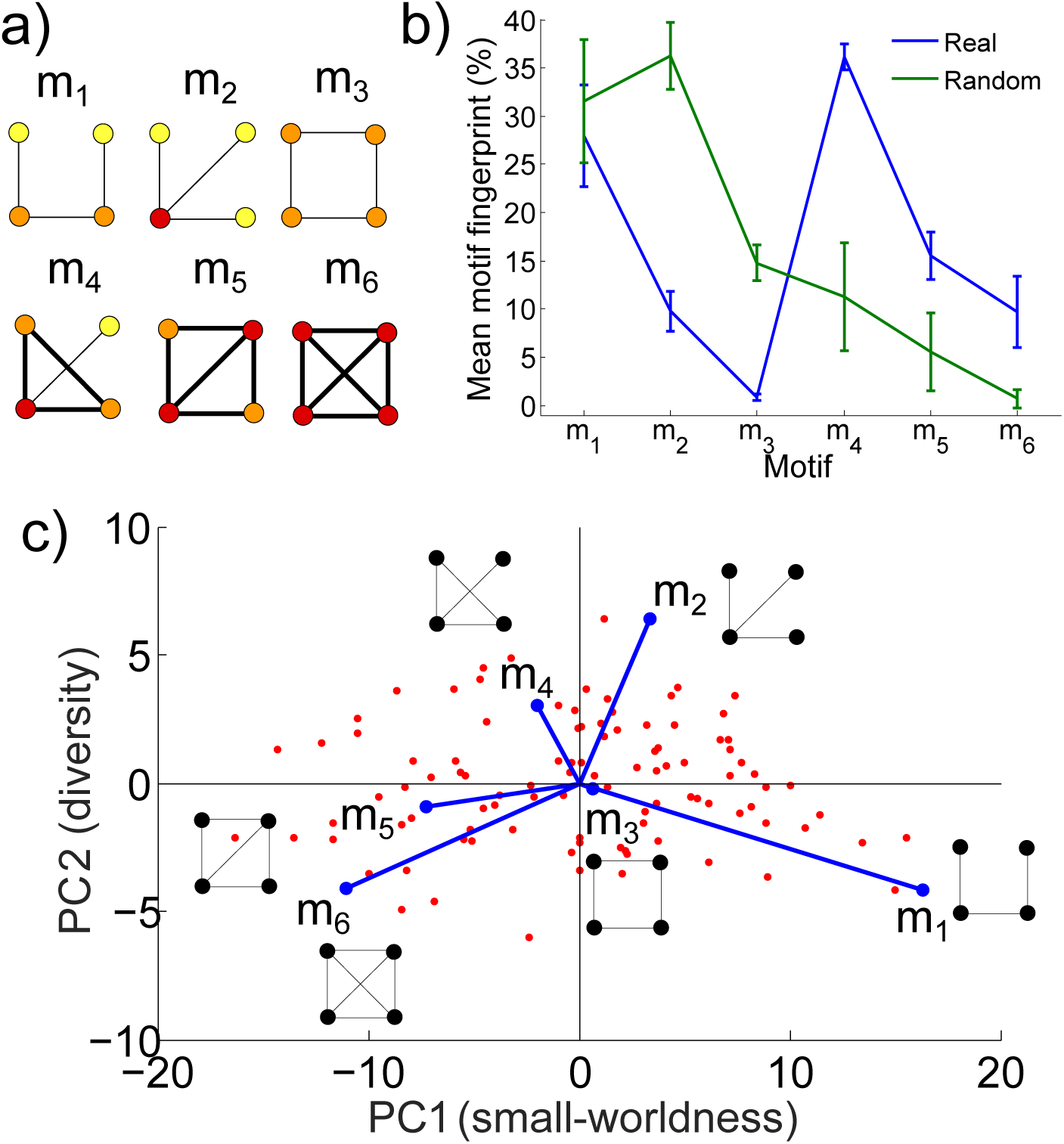
Motifs, motif fingerprint and motif morphospace. a) All six possible 4-node undirected motifs. Motif nodes are coloured according to their degree, where yellow nodes have degree 1, orange have degree 2 and red have degree 3. Motif edges forming triangles are highlighted in bold. b) Motif fingerprint averaged across all subjects (blue). The error bars show ±*σ*, where *σ* is the standard deviation. The motif fingerprint from randomised networks are shown in green for comparison (the degree distribution was preserved and results were averaged over 100 realisations.) c) The motif PC morphospace plotted using the first two PCs. Here each red dot represents an individual subject, plotted in PC space. The original motifs are also plotted as vectors in the PC space. The axes are labelled as small-worldness and diversity due to the correlations of PC1 and PC2 with global network metrics, see below.

For each individual network, we count the number of each of these motifs and normalise with respect to the total number of motifs in the network. This gives us a motif ‘fingerprint’ for each network; further details are given in the Methods section. The average fingerprint across all networks is shown in Fig. 1b, alongside the standard deviation across the subjects and the motif fingerprint for randomised networks for comparison. We observe a high proportion of *m*_1_ and *m*_4_. Motif *m*_3_ has the lowest proportion, which is expected because the networks are obtained by calculating the correlations between time series, hence they exhibit a high proportion of closed triangles [33]. We observe intermediate proportions of *m*_2_*, m*_5_ and *m*_6_.

We perform a PCA on the motif fingerprints for all 100 subjects, again as described in the Methods section. The first PC explains 86% of the total variability and the first 2 PCs explain 97% of the total variability; see the cumulative variability plot shown in Supplementary Figure 1. Note that here we use 4-node motifs, however similar results can be found using the 21 possible 5-node motifs (see section 6 of the SI). Hence we focus on the first two motif PCs. By plotting the individual networks as a function of these PCs, we can create a two dimensional morphospace, as shown in Fig. 1c (here individual subjects are represented by red dots). In order to visualise the extent to which different motifs underlie the PCs, we also plot the original motif fingerprint variables as vectors in the PC space (sometimes known as a ‘biplot’). We find that PC1 is driven by a difference between the ‘chain-like’ *m*_1_ and the more densely clustered *m*_5_ and *m*_6_. Networks with high PC2 values have a high proportion of *m*_2_ and *m*_4_, which have greater variability in motif node degree than the other motifs. Remarkably, using 5-node motifs produces a very similar biplot to the 4-node motifs, with a similar pattern of motifs underlying the morphospace, as shown Supplementary Figure 7b. We used a threshold of 800 edges (≈ 20% connection density), however similar motif fingerprints and biplots were obtained with 400 and 1200 edges (≈ 10% or 30% connection density), see section 2 of the SI. We also reproduced the results using a different parcellation (the AICHA parcellation [12], with 384 regions rather than 90), as well as in the HCP re-test dataset and in an independent dataset, see sections 3, 4 and 5 of the SI, respectively.

### Relationship between the PCs and global network metrics

To investigate the differences in the networks which underlie the PC space further, we calculate the Pearson correlations between the PCs and global network metrics, namely global efficiency, assortativity and transitivity. The correlations are summarised in Fig. 2a and in Fig. 2b we plot the PC space coloured according to the global efficiency and assortativity. With the exception of a weak correlation between PC 3 and global efficiency (with Pearson correlation coefficient r=-0.22, and a p-value for a two-tailed Pearson correlation test using the student’s t-distribution of p=0.03), PCs 3, 4, 5 and 6 did not correlate significantly (p> 0.05) with any of the global network measures tested. This is as expected because 97% of the motif variability is explained by PCs 1 and 2.

**Figure 2.**
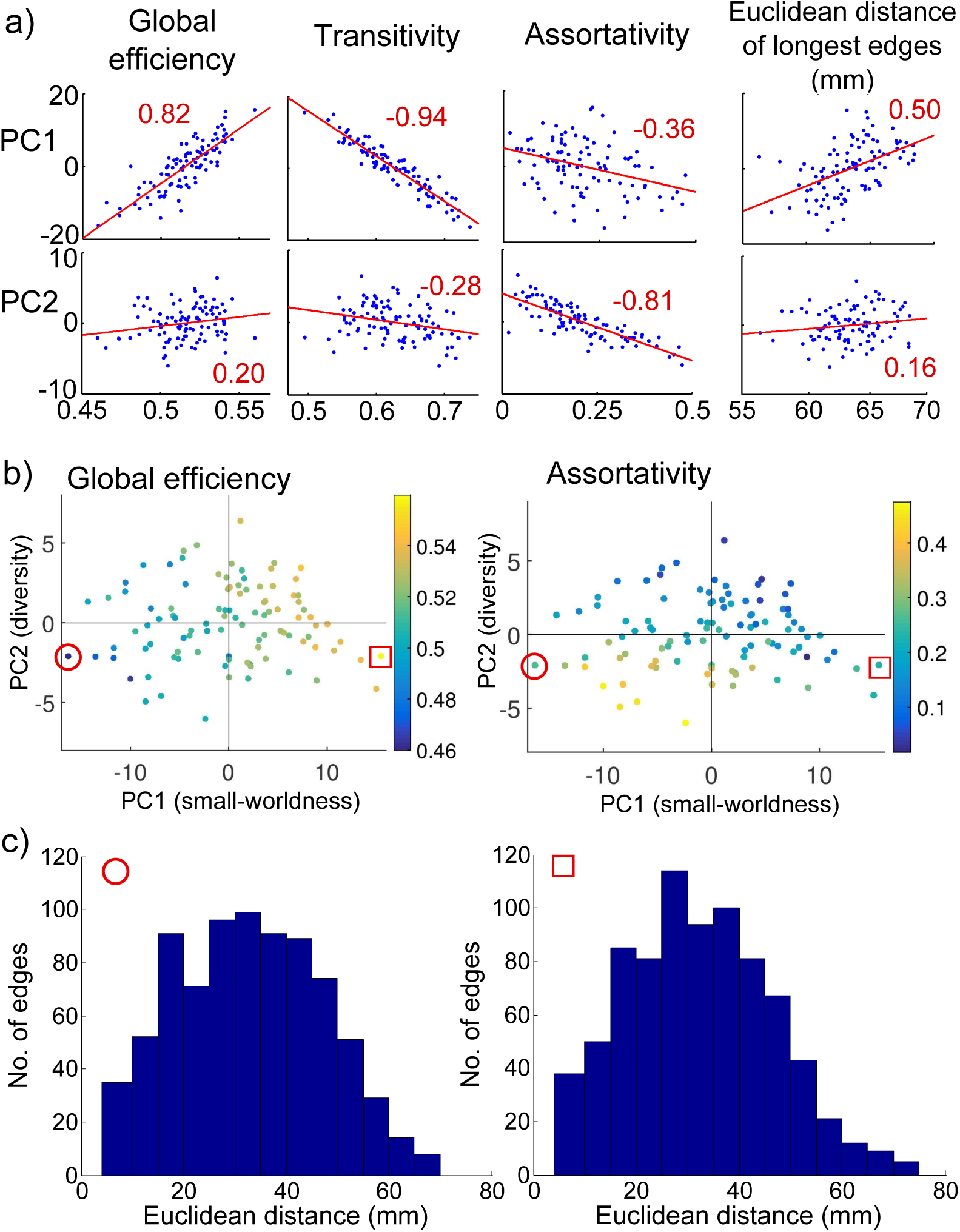
Motif morphospace dimensions and global topology. a) The correlation between the first two PCs and the global efficiency, assortativity and transitivity of the networks. The correlation between the PCs and the average Euclidean distance of the longest 5% of edges is also shown. In all plots, the Pearson correlation coefficient is shown in red. b) The motif morphospace (as shown in Fig. 1c), coloured according to the global efficiency and the assortativity of each network. c) Edge distance distributions for a subject with high PC1 (denoted by a red square) and a subject with low PC2 (denoted by a red circle).

PC1 is positively correlated with global efficiency (with a Pearson correlation coefficient r=0.82, p< 0.001) and negatively correlated with transitivity (r=-0.94, p< 0.001). Hence as the proportion of the ‘chain-like’ *m*_1_ increases and the proportion of the fully clustered *m*_6_ decreases, the global efficiency of the networks tends to increase, whilst the transitivity decreases. This inverse relationship between global efficiency and transitivity is reminiscent of the trade-off navigated by small-world networks, which can exhibit both high clustering and high global efficiency [31]. fMRI brain networks are known to exhibit small-world properties [3, 4] and our results provide further evidence that this is a key axis of variability in their topologies. We note that in 2012 [27] suggested a heuristic model of brain networks (Fig. 10 in their paper) in which one of the main axes is defined by small-worldness and is similar to our first PC. Hence we label PC1 as ‘small-worldness’.

PC2 is most strongly correlated with assortativity (r=-0.81, p< 0.001), hence networks with higher proportions of *m*_2_ and *m*_4_ tend to have lower assortativity values. The observation that apical motifs (e.g. *m*_2_) tend to have high PC2 scores suggests that PC2 might also be expected to correlate with hierarchy. The measure of hierarchy proposed by Mones et al. [20], which measures the heterogeneity in closeness centrality of the network’s nodes, does correlate strongly with PC2 (r=0.74, p< 0.001) and more weakly with PC1 (r=-0.41, p< 0.001). Ravasz et al. studied hierarchical modularity using the scaling of the nodes’ clustering coefficients with their degrees [23]. The exponent of this relationship correlates most strongly with PC1 (r=0.63, p< 0.001) and less strongly with PC2 (r=0.40, p< 0.001). This difference reflects the fact that Ravasz’s approach is more sensitive to the networks’ modularity than Mones’ measure. PC2 is not dissimilar to Stam’s vertical axis, which he labels ‘diversity’ [27]. We note that from Fig. 1c, motifs which show high PC2 values tend to have a higher level of node degree diversity than motifs with low PC2 values. Therefore we label PC2 as ‘diversity’.

We note that the correlation of PC1 with the efficiency and transitivity of the networks and PC2 with the assortativity described above are extremely robust. We found that they can be replicated with 400 or 1200 edges instead of 800, as shown in section 2 of the SI. The results were also robust to changing the parcellation to the AICHA parcellation [12], which has 384 regions rather than 90, see section 3 of the SI. Lastly, the results were reproduced both in the re-test HCP dataset and the independent ‘Cambridge’ dataset described in the Methods section. For details, see sections 4 and 5 of the SI.

### Relationship between the PCs and the spatial embedding of the networks

Having observed significant correlations between the PCs and global network metrics, we now turn to the relationship between the PCs and the spatial embedding of the networks. We take Euclidean distance as a measure characterising the spatial embedding. We begin by calculating the correlation between PCs 1 and 2 and the total Euclidean distance of each network’s connections and obtain r=0.03 (p=0.75) and r=0.17 (p=0.09) respectively. We conclude that there is no significant correlation between total Euclidean distance and PCs 1 and 2. Note that there is also no significant correlation (p > 0.05) between the global efficiency, assortativity or transitivity and the total Euclidean distance.

However, whilst there is no correlation between the PCs and the total Euclidean distance, we do observe significant correlation between PC1 and the average length of the longest 5% of edges in each network, with r=0.50 (p< 0.001), as shown in Fig. 2a. This positive correlation means that the longest 5% of edges tend to be longer in networks with a relatively high proportion of ‘chain-like’ *m*_1_ (which from Fig. 1 also tend to be those with higher global efficiency). In Fig. 2c, we plot the distance distribution for the subject with the highest and lowest PC1 values. As expected, the distance distribution for the subject with a high PC1 value has a longer tail at long distances. In contrast, there is no significant correlation between the length of the longest 5% of edges and PC2 (r=0.16, p=0.11). The significant correlation between PC1 and the length of the longest edges is robust to changing the percentage of the longest edges considered from 1% to 10% and to considering 400 or 1200 edges instead of 800 (see sections 2 and 8 of the SI). We also reproduced the result in the re-test HCP dataset and obtained r=0.43 (p< 0.001). Interestingly, the result was not significant (p > 0.05) in an independent dataset with 26 subjects (the ‘Cambridge’ dataset) as discussed in section 5 of the SI. This may be due to the reduced number of subjects available, however further work is required to clarify the reason for this difference.

### Comparison with generative models

The motif morphospace described above enables us to map the global topology of fMRI brain networks and to explore the relationships between different global properties of the networks. Previous work has suggested that a simple two parameter generative model can capture many of the global features of fMRI brain networks [30]. Hence in the final part of this work we investigate the extent to which a simple two parameter generative model can capture the motif morphospace we propose.

Vértes et al. found that the economical clustering generative model given by equation (1) was able to reproduce many aspects of fMRI brain network topology. This generative model encapsulates the trade-off between the cost of long distance connections and the network’s topology. Figure 3a illustrates the factors which determine the probability of connecting nodes *i* and *j* (*P_i_,_j_*) in the model. The length of the connection between nodes *i* and *j* is approximated by their Euclidean distance, *d_i,j_* and the probability of connecting the nodes depends on this distance and the parameter *η.* The topological term employed by the model sets the probability of connecting nodes *i* and *j* to increase with the number of nearest neighbours they have in common, *k_i,j_.* The strength of this dependence is given by the parameter *γ.* Overall, *P_i,j_* is given by:

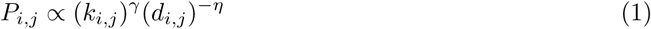

**Figure 3.**
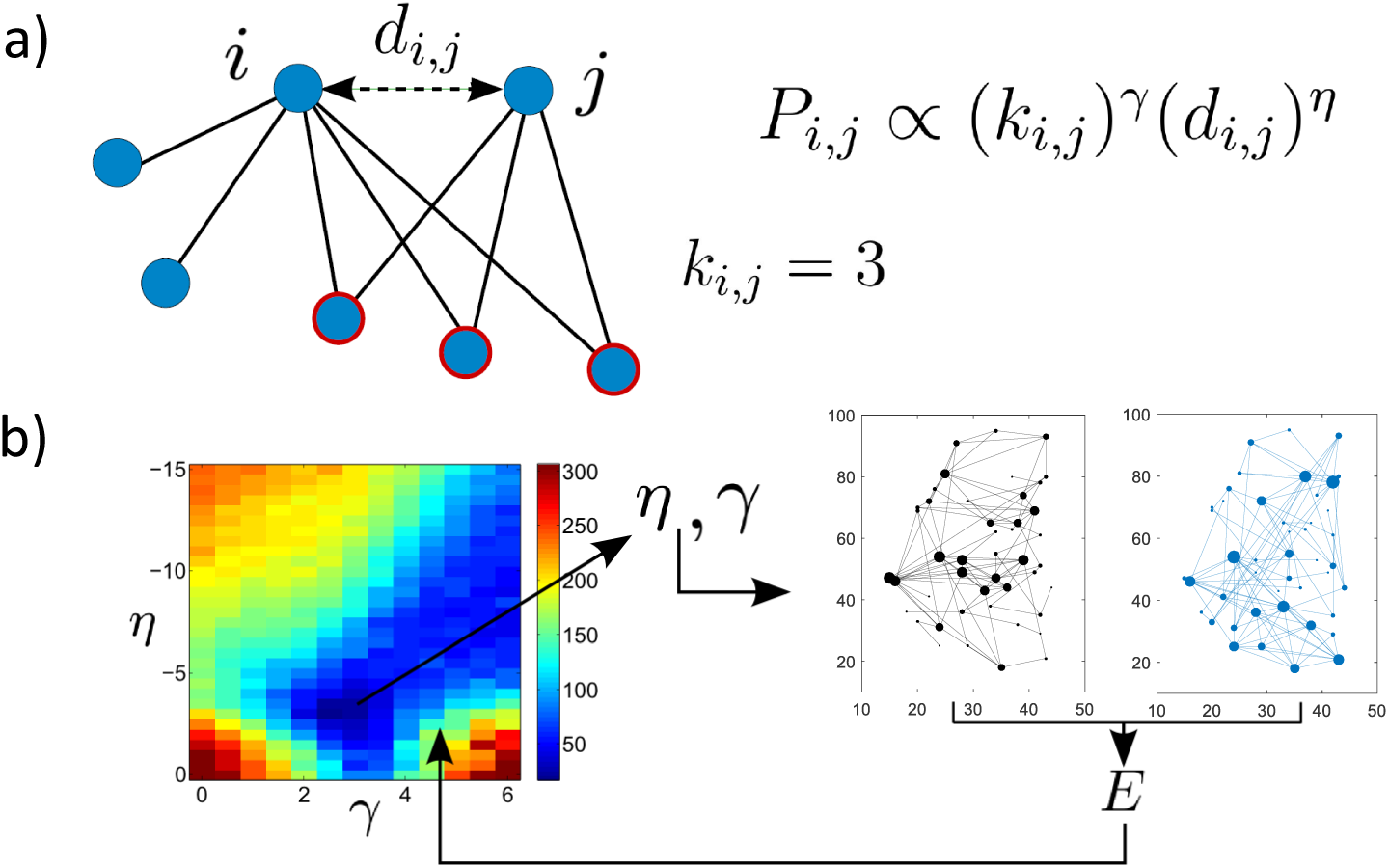
Schematic of economical clustering generative model. a) The probability of a new link being added between nodes *i* and *j* depends on both the Euclidean distance between them (d_i,j_) and the number of nearest neighbours they have in common (k_i,j_ = 3 in this example). b) Illustration of the generative model steps. Networks are generated at a range of different *η* and *γ* values and their fit to the real brain networks is calculated via an energy function, E, which compares networks on the basis of global efficiency, modularity, transitivity and degree distribution (as used by [30] et al.). On the left hand side the energy is plotted as a function of *η* and *γ.* On the right hand side, an example real brain network and generated network are shown for illustration (in black and blue, respectively). The final generated networks are simulated using the optimised values of *η* and *γ.*

We use the economical clustering model to simulate the topology of our fMRI networks. All of the results described up to this point are robust to thresholding the networks at either 400, 800 or 1200 edges (≈ 10%, 20% or 30% connection density). However, the generative models require relatively sparse networks, therefore we use a threshold of 400 edges in this section. We also use a single hemisphere (left) rather than both hemispheres, as in [30] and [5].

We use an independent dataset and an identical approach to generating the networks as [30]. We determine the parameters (*γ* and *η*) by exploring the parameter space manually to obtain values of *γ* and *η* which optimise the global efficiency, modularity, mean clustering coefficient and degree distribution of the generated networks with respect to the real HCP fMRI brain networks. For our networks, which have fewer nodes than those used in [30], we find that *η* = −3 and *γ* = 2.5 are the optimal values, see section 8.1 of the SI for more details. Note that whilst these parameters were optimised for the measures listed above, they were not optimised to match the motif fingerprints or other network measures including the assortativity. We stress that the real fMRI networks were only used to optimise *η* and *γ* and the simulated networks were generated with no other knowledge of the real networks. The process is illustrated in Fig. 3b.

In Fig. 4a, we plot the average motif fingerprint of the generated networks alongside the real fMRI brain networks and note that they are the same to within ±*σ*, the standard deviation. When we project the generated networks onto the original morphospace of fMRI brain networks, we find that they lie in a similar region, see Fig. 4b. In Fig. 4c, we plot the biplot of the generated networks, which is remarkably similar to the biplot shown in Fig. 1c. The correlation between the PCs and the global efficiency, assortativity and transitivity are similar to those obtained for the real fMRI brain networks; see the SI for details. The correlation between the average Euclidean distance of the longest 5% of edges and PC1 is not statistically significant (r=0.18 and p=0.07). This small correlation is partly, although not solely, because only a single hemisphere was considered, hence there are fewer long distance connections. Note that other two parameter models, for example the economical preferential attachment generative model also discussed by Vértes et al., are not able to reproduce the motif fingerprint of the real fMRI brain networks, as demonstrated in section 8.2 of the SI.

**Figure 4.**
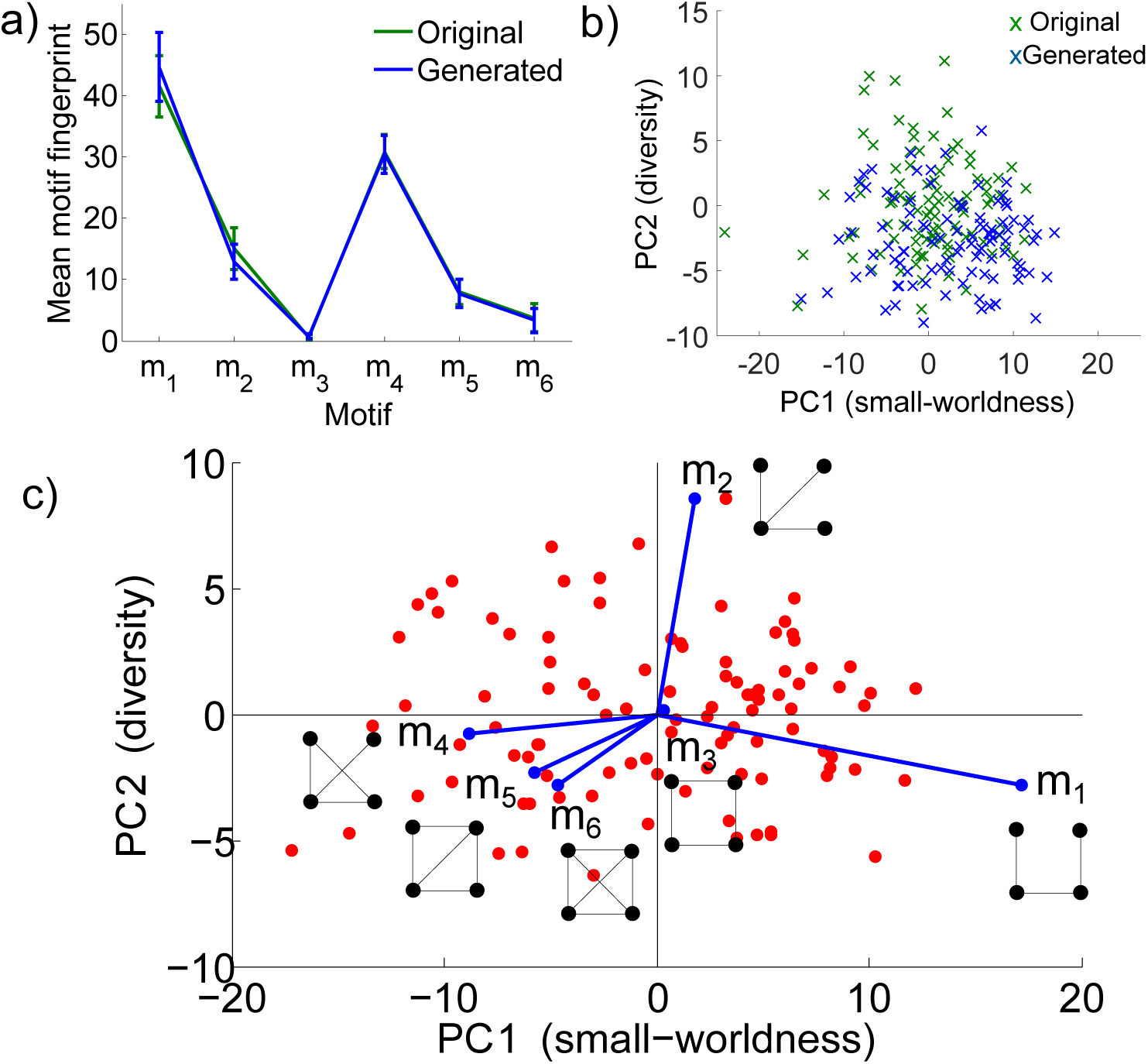
Generative model results. a) The motif fingerprints of the original real brain networks (with a single hemisphere and 400 edges), alongside the motif fingerprints of the generated networks (*η* = −3 and *γ* = 2.5). b) The generated networks projected onto the space of the original networks. c) The biplot of the generated networks.

One advantage of the morphospace framework is that it allows us to explore the effect of changing the generative model parameters and to explore how the factors of the model relate to one another. Therefore, lastly we generate networks with different *η,* setting *γ* = 0 and vice versa, and project the results onto the morphospace of our original fMRI brain networks. Note that the axes of the morphospace are still defined by the real fMRI brain networks. The results are shown in Fig. 5. We note that changing either *η* or *γ* can change the networks’ positions along both PC1 and PC2, suggesting that both the distance and clustering penalisations can influence the networks’ motif fingerprints in similar ways. The heat maps of global network properties with different *η* and *γ* values shown in Supplementary Figure 8 also show that the distance and clustering parameters can compensate for each other to some extent.

**Figure 5.**
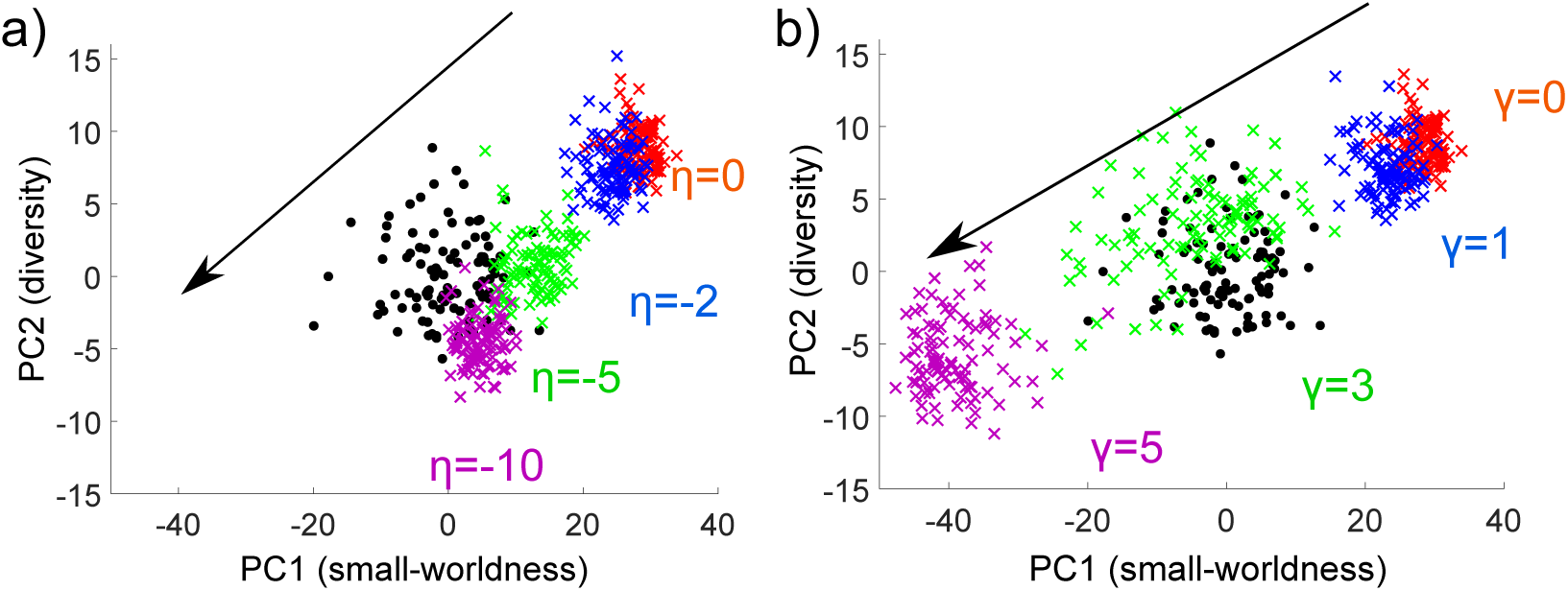
Generated networks with different *η* and *γ*. a) Generated networks with different *η* and *γ* = 0 projected onto the space of the original networks. b) Generated networks with different *γ* and *η* = 0 projected onto the space of the original networks. In both parts the original networks are shown as black dots and the superimposed networks are shown as coloured crosses. Arrows denote the direction of increasing |*η*| or *γ.*

## Discussion

Our results show that, surprisingly, 97% of the variability in the motif fingerprints of fMRI brain networks can be explained by just two PCs. The fact that these data driven PCs correlate significantly with several global metrics, including global efficiency and assortativity, suggests that these manually defined metrics do capture key variations within the data. This alignment between manually defined and data driven measures is non-trivial and gives confidence in the graph metrics. Our motif morphospace allows us to go a step further and to map the global topology of the networks more comprehensively than any individual network measure. The morphospace also enables us to explore the extent to which different metrics overlap and the relationships between them. Crucially, the space we obtain is low-dimensional, suggesting that the global topology of fMRI brain connectivity networks can be explained by few parameters.

The PCs of the morphospace represent the two main degrees of variability in the networks. The first PC (which alone explains over 80% of the variability) is positively correlated with global efficiency and the average length of the longest 5% of edges and negatively correlated with transitivity. Hence the main axis of variability in the structure of healthy fMRI networks represents the difference between networks with high numbers of long distance edges and high global efficiency, versus those with fewer long distance edges, low global efficiency and high transitivity. This can be thought of as the small-world axis, in line with many reports in the literature that small-worldness is a key factor in fMRI brain networks and encodes a trade-off between the topological benefits of long distance edges and their cost [3, 4, 27]. PC2 is more novel and correlates most strongly with the assortativity of the networks. Assortativity has been linked to network robustness in the literature [21], however further work is needed to understand the exact role of assortativity in fMRI brain networks.

Generative models allow us to shed further light on the driving forces behind the two PCs. The motif fingerprints of the networks and their motif biplot can be reproduced using the two parameter economical clustering generative model proposed by [30]. Other generative models - including the economical preferential attachment model- do not reproduce the space as well. This suggests that the nearest neighbour model is closer to the underlying biological mechanism than the preferential attachment model. This observation also demonstrates that it is not trivial to obtain a two parameter model which can reproduce the networks (despite optimising the model parameters) and shows that the motif morphospace provides a powerful way to distinguish between different generative models. In future, we propose that models trying to reproduce fMRI brain networks use this motif morphospace approach to test their validity and to compare different models.

Varying the parameters *η* or *γ* in the generative model and projecting the resulting networks into the real fMRI brain network morphospace changes the networks’ motif fingerprints and their positions along both PC1 and PC2. This suggests that both *η* and *γ* are important in determining the networks’ motif fingerprints and positions along both PC1 and PC2 of the morphospace. Note that it is important to distinguish between:

1. the variation in the positions of networks which all have a set value of *η* and *γ,* and
2. the variation in the positions of superimposed networks with different *η* and *γ* parameter values.

The former is due to random variations in the edges which are created *(η* and *γ* only set the probability of creating edges), whilst the latter is due to the changes in *η* and *γ.* One interesting question is what causes the topological differences in healthy fMRI brain networks described by the two PCs in our motif morphospace. Our results show that we can model a large amount of the variability in healthy fMRI networks with random fluctuations in a generative model. This suggests that the healthy population all follow the same underlying trade-offs (the parameters in the generative models), with fluctuations in how those trade-offs are navigated. For example, healthy brains make the same small-worldness trade-off between the topological benefits of long distance edges and their cost, with fluctuations in the exact compromise reached. Unlike in healthy subjects, [30] suggested that in diseased states such as schizophrenia the underlying generative rules themselves are altered. Further work is required to clarify the microscopic basis underlying these generative rules and their relationship with brain function.

Overall, the motif morphospace provides a data driven, simple way to study and visualise network topology in a low-dimensional space. Our results suggest that this approach could also be useful in characterising other types of networks. Extending the approach to directed networks is straightforward. We note that in principle a motif morphospace could also be used to compare different types of networks.

## Methods

We use 100 fMRI brain scans of healthy individuals from the Human Connectome Project. These data were already used in [28] to assess the reproducibility of the graph metrics. In this context, we downloaded the resting state fMRI dataset publicly released as part of the Human Connectome Project (HCP), WU-Minn Consortium (for detailed parameters see [25]). The functional images were acquired on a 3T Siemens Connectome Skyra MRI scanner with a 32-channel head coil, using a multiband gradient-echo EPI imaging sequence with the following parameters: 2 mm isotropic voxels, 72 axial slices, TR = 720 ms, TE = 33.1 ms, flip angle = 52, field of view = 208x180 mm^2,^ matrix size = 104x90 and a multiband factor of 8. A total of 1200 images was acquired for a scan duration of 14 min and 24 s. The data used from the HCP include a second dataset with the same 100 subjects scanned a second time on different days with identical acquisition parameters. This re-test dataset allowed us to confirm and validate our results.

In order to extract brain connectivity graphs, the structural and functional data were preprocessed according to the pipeline described by [8]. Finally, the functional data were registered to the individual structural image and further to the MNI152 atlas space using the transforms applied to the structural image (see [28] for details). In the present study, each brain image is parcellated using the classical Anatomic-Automatic Labeling (AAL) [29] composed of 90 regions. The results were also robust to using a different parcellation (the AICHA parcellation, with 384 regions), as shown in section 3 of the SI.

In each parcel, regional mean time series are estimated by averaging, at each time point, the fMRI voxel values weighted by the grey matter probability of these voxels. Correlation matrices with 90×90 elements are computed using wavelets [1], and wavelet scale four (corresponding to frequency interval [0.043 – 0.087] Hz) is chosen for extracting the graphs by thresholding the correlation matrices. The correlation threshold is applied to each individual network so that all graphs have the same number of edges. The cost, that is the ratio between the number of edges in the graph and the total number of possible edges, is fixed to approximately 20% (800 edges) based on the reproducibility results obtained in [28]. Indeed, it was shown that for 20% cost, the test-retest reproducibility is higher for the majority of graph metrics (in this work we also test other thresholds, as shown in the SI). Each subject brain connectivity network is then represented as a graph G = (V, E) where V is the number of vertices or parcels (90 in our study), and E is the number of edges (800).

We then turn to characterise the decomposition of each graph G into motifs. We use the software ‘FANMOD’ to count the number of motifs in each of the 100 networks [32] (other software can also be used, for example ‘mfinder’ [19] or ‘orca’ [11]). Unless otherwise stated, we focus on the six possible 4-node undirected induced motifs shown in Fig. 1a, which we denote as *m_i_,* where *i* ∈ 1, 6. We count the number of each of these motifs within each network, *N_i_*, and then normalise the number of motif i with respect to the total number of motifs in the network, to obtain the motif fingerprint for each network: 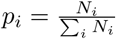.

We then perform a principal component analysis on the motif fingerprints for all of the 100 HCP networks. Here we have 100 observations (the subjects) and 6 variables (the fingerprint scores for each of the six possible 4-node undirected induced motifs). The PCA was performed in MATLAB. The results of the PCA are displayed using a biplot of the principal component coefficients. The axes of the biplot represent the principal components, the observations are shown as red dots and the original variables are plotted in the PC space as vectors.

The global efficiency, assortativity, modularity and transitivity of the networks were calculated using the Brain Connectivity Toolbox [24]. Note that transitivity is calculated as the ratio of triangles to triplets in the network (sometimes known as clustering). Modularity is calculated as the maximised modularity Q obtained by Newman’s spectral community detection [22]. The total Euclidean distance is a sum of the Euclidean distance of all edges in the thresholded network.

The generative models we compare the brain networks to are generated using the economical clustering model, given by equation (1). We explore the parameter space manually to determine the values of *η* and *γ* which optimise the global efficiency, mean clustering coefficient, modularity and degree distribution of the real HCP fMRI brain networks (see the SI for more details). Following [30], we generate the networks with both hemispheres, but only consider a single hemisphere in subsequent calculations since the links between hemispheres are expected to follow different patterns to those within a single hemisphere. We use a threshold of 400 edges, as discussed in the Results section, and include a small number of additional edges if necessary to ensure that the graphs are connected.

## Supporting Information

Supplementary information is attached. Code is available online at: https://github.com/SarahMorgan/Motif-Morphospace.

## Acknowledgments

P.E.V. is supported by a Medical Research Council Bioinformatics Research Fellowship (grant number MR/K020706/1). M.T. was supported by a grant from the Rhône-Alpes Région, France. S.A. was partly funded by a grant from la Région Rhône-Alpes and a grant from AGIR-PEPS, Université Grenoble Alpes–CNRS. Data were provided by the Human Connectome Project, WU-Minn Consortium (Principal Investigators: David Van Essen and Kamil Ugurbil; 1U54MH091657) funded by the 16 NIH Institutes and Centers that support the NIH Blueprint for Neuroscience Research; and by the McDonnell Center for Systems Neuroscience at Washington University. The authors are grateful to Prof. Uri Alon, Dr Miri Adler and Dr Pablo Szekely for sharing code.

## Author Contributions

S.E.M, P.E.V and E.T.B designed the study, S.E.M. performed the simulations and data analysis. S.A. and M.T. pre-processed the data. S.E.M., S.A., E.T.B. and P.E.V. discussed the results and wrote the paper.

## References

1. S. Achard, R. Salvador, B. Whitcher, J. Suckling, and E. Bullmore. A resilient, low-frequency, small-world human brain functional network with highly connected association cortical hubs. The Journal of Neuroscience, 26(1):63–72, Jan. 2006. 00831.

2. A. Avena-Koenigsberger, J. Goñi, R. Solé, and O. Sporns. Network morphospace. Journal of the Royal Society Interface, 12:20140881, 2014.

3. D. Bassett and E. Bullmore. Small-world brain networks revisited. Neuroscientist, 12:512–23, 2006.

4. D. Bassett and E. Bullmore. Small-world brain networks. Neuroscientist, page 1073858416667720,2016.

5. R. Betzel, A. Avena-Koenigsberger, J. Goñi, Y. He, M. de Reus, A. Griffa, P. Vértes, B. Mišic, J. Thiran, P. Hagmann, M. van den Heuvel, X. Z. Zuo, E. Bullmore, and O. Sporns. Generative models of the human connectome. NeuroImage, 124:1054–1064, 2016.

6. E. Bullmore and O. Sporns. The economy of brain network organization. Nature Reviews Neuroscience, 13:336–349, 2012.

7. R. Diestel. Graph Theory. Springer, 2006.

8. M. F. Glasser, S. N. Sotiropoulos, J. A. Wilson, T. S. Coalson, B. Fischl, J. L. Andersson, J. Xu, S. Jbabdi, M. Webster, J. R. Polimeni, D. C. Van Essen, and M. Jenkinson. The minimal preprocessing pipelines for the human connectome project. NeuroImage, 80:105–124, Oct. 2013.

9. J. Goñi, A. Avena-Koenigsberger, N. V. de Mendizabal, M. P. van den Heuvel, R. F. Betzel, and O. Sporns. Exploring the morphospace of communication efficiency in complex networks. PLOS ONE, 8:e58070, 2014.

10. J. Henderson and P. Robinson. Geometric effects on complex network structure in the cortex. Physical Review Letters, 107:018102, 2011.

11. T. Hočevar and J. Demšar. A combinatorial approach to graphlet counting. Bioinformatics, 30:559–565, 2014.

12. M. Joliot, G. Jobard, M. Naveau, N. Delcroix, L. Petit, L. Zago, F. Crivello, E. Mellet, B. Mazoyer, and N. Tzourio-Mazoyer. Aicha: An atlas of intrinsic connectivity of homotopic areas. J. Neurosci. Methods, 254:46–59, 2015.

13. M. Kaiser and C. Hilgetag. Modelling the development of cortical networks. Neurocomputing, 58:297–302, 2004.

14. M. Kaiser and C. Hilgetag. Spatial growth of real world networks. Physical Review E, 69:036103, 2004.

15. M. Kaiser and C. C. Hilgetag. Nonoptimal component placement, but short processing paths, due to long-distance projections in neural systems. PLOS Computational Biology, 2(7):e95, 2006.

16. F. Klimm, D. Bassett, J. Carlson, and P. Mucha. Resolving structural variability in network models and the brain. PLOS Computational Biology, 10:e1003491, 2013.

17. C. Li, H. Wang, W. de Haan, C. J. Stam, and P. Van Mieghem. The correlation of metrics in complex networks with applications in functional brain networks. Journal of Statistical Mechanics: Theory and Experiment, 11:11018, 2011.

18. R. Milo, S. Itzkovitz, N. Kashtan, R. Levitt, S. Shen-Orr, I. Ayzenshtat, M. Sheffer, and U. Alon. Superfamilies of evolved and designed networks. Science, 303:1538–1542, 2004.

19. R. Milo, S. Shen-Orr, S. Itzkovitz, N. Kashtan, D. Chklovskii, and U. Alon. Network motifs: Simple building blocks of complex networks. Science, 298:824–827, 2002.

20. E. Mones, L. Vicsek, and T. Vicsek. Hierarchy measure for complex networks. PLOS ONE, 7:e33799, 2012.

21. M. Newman. Assortative mixing in networks. Phys. Rev. Lett., 89:208701, 2002.

22. M. Newman. Finding community structure in networks using the eigenvectors of matrices. Phys. Rev. E, 74:036104, 2006.

23. E. Ravasz and A.-L. Barabasi. Hierarchical organization in complex networks. Physical Review E, 67:026112, 2003.

24. M. Rubinov and O. Sporns. Complex network measures of brain connectivity: Uses and interpretations. NeuroImage, 52:1059–69, 2010.

25. S. M. Smith, C. F. Beckmann, J. Andersson, E. J. Auerbach, J. Bijsterbosch, G. Douaud, E. Duff, D. A. Feinberg, L. Griffanti, M. P. Harms, M. Kelly, T. Laumann, K. L. Miller, S. Moeller, S. Petersen, J. Power, G. Salimi-Khorshidi, A. Z. Snyder, A. T. Vu, M. W. Woolrich, J. Xu, E. Yacoub, K. Uǧurbil, D. C. Van Essen, and M. F. Glasser. Resting-state fMRI in the human connectome project. NeuroImage, 80:144–168, Oct. 2013.

26. O. Sporns and R. Kötter. Motifs in brain networks. PLOS Biology, 2(11), 10 2004.

27. C. Stam and E. van Straaten. The organization of physiological brain networks. Clinical Neurophysiology, 123:1067–1087, 2012.

28. M. Termenon, A. Jaillard, C. Delon-Martin, and S. Achard. Reliability of graph analysis of resting state fMRI using test-retest dataset from the human connectome project. NeuroImage, 142:172–187, 2016.

29. N. Tzourio-Mazoyer, B. Landeau, D. Papathanassiou, F. Crivello, O. Etard, N. Delcroix, B. Mazoyer, and M. Joliot. Automated anatomical labeling of activations in SPM using a macroscopic anatomical parcellation of the MNI MRI single-subject brain. NeuroImage, 15(1):273–289, Jan. 2002.

30. P. Vértes, A. Alexander-Bloch, N. Gogtay, J. Giedd, J. Rapoport, and E. Bullmore. Simple models of human brain functional networks. Proc. Natl. Acad. Sci., 109:5868-73, 2012.

31. D. J. Watts and S. H. Strogatz. Collective dynamics of ‘small-world’networks. nature, 393(6684):440-442, 1998.

32. S. Wernicke and F. Rasche. FANMOD: a tool for fast network motif detection. Bioinformatics, 22:1152-1153, 2006.

33. A. Zalesky, A. Fornito, and E. Bullmore. On the use of correlation as a measure of network connectivity. NeuroImage, 60:2096-2106, 2012.

